# The plastoglobule-localized AtABC1K6 is a Mn^2+^-dependent protein kinase necessary for timely transition to reproductive growth

**DOI:** 10.1101/2021.10.28.466314

**Authors:** Roberto Espinoza-Corral, Peter K. Lundquist

## Abstract

The <u>A</u>bsence of <u>bc_1_</u> Complex (ABC1) is an ancient, atypical protein kinase family that emerged prior to the archaeal-eubacterial divergence. Loss-of-function mutants in ABC1 genes are linked to respiratory defects in microbes and humans, and to compromised photosynthetic performance and stress tolerance in plants. However, demonstration of protein kinase activity remains elusive, hampering their study. Here, we investigate a homolog from *Arabidopsis thaliana*, AtABC1K6, and demonstrate *in vitro* protein kinase activity as autophosphorylation, which we replicate with a human ABC1 ortholog. We show that AtABC1K6 protein kinase activity requires an atypical buffer composition, including Mn^2+^ as divalent cation co-factor and a low salt concentration. AtABC1K6 associates with plastoglobule lipid droplets of *A. thaliana* chloroplasts, along with five paralogs. Protein kinase activity associated with isolated *A. thaliana* plastoglobules was inhibited at higher salt concentrations, but could accommodate Mg^2+^ as well as Mn^2+^, indicating salt sensitivity, but not the requirement for Mn^2+^, may be a general characteristic of ABC1s. Loss of functional AtABC1K6 impairs the developmental transition from vegetative to reproductive growth. This phenotype is complemented by the wild-type sequence of AtABC1K6 but not by a kinase-dead point mutant in the unique Ala-triad of the ATP-binding pocket, demonstrating the physiological relevance of the protein’s kinase activity. We suggest that ABC1s are bona fide protein kinases with a unique regulatory mechanism. Our results open the door to detailed functional and mechanistic studies of ABC1s and plastoglobules.

**SIGNIFICANCE STATEMENT:** The <u>A</u>bsence of <u>bc_1_</u> Complex (ABC1) is an ancient, atypical protein kinase family with enigmatic physiological roles in a wide range of species including plants, humans and microbes. While mutants demonstrate their critical role for organismal survival, their study has been severely hampered by the previous inability to determine catalytic function. Here, we demonstrate *in vitro* protein kinase activity with an *A. thaliana* homolog, AtABC1K6. Loss of functional AtABC1K6 impairs the developmental transition from vegetative to reproductive growth. The lack of phenotypic complementation with a kinase-dead point mutant demonstrates the physiological relevance of the protein’s kinase activity. Our results present the experimental means to investigate the targets, functions, and regulation of ABC1s.

## INTRODUCTION

Plastoglobule lipid droplets are a unique compartment of plant chloroplasts that are physically associated with the photosynthetically active thylakoid membrane (1). They hold essential but poorly understood roles related to photosynthesis, development, and stress tolerance (2–4). Despite the persistent physical association, the plastoglobule maintains a proteome and lipidome that is distinct from the thylakoid, comprising a rich store of various prenyl-lipid compounds related to photosynthesis (i.e. tocochromanols, quinones, and carotenoids), and several dozen proteins largely related to prenyl-lipid metabolism or redox reactions (5, 6). We have previously proposed that the plastoglobule serves as a dynamic hub that orchestrates rapid thylakoid membrane remodeling, sequesters and neutralizes toxic (oxidized) lipid and protein, and scavenges reactive oxygen species (5, 6). These functions are thought to be engaged routinely to regulate photosynthesis under constantly changing environmental conditions and to promote developmental transitions. This model presumes rapid functional modulation of the plastoglobule proteome, however, it remains unclear how such rapid modulation may be effected.

Prime candidates to achieve this modulation are six members of an ancient, atypical protein kinase (aPK) family known as the <u>a</u>ctivity of <u>bc_1_</u> complex (ABC1) family (alternately known as COQ8, UbiB, or aarF domain-containing kinase [ADCK]) (7). ABC1s are conserved among archaea, eubacteria, and eukaryotes where mutant phenotypes have indicated a critical role in ubiquinone biosynthesis (7–9), reflected in the various names ascribed to the family. While one to two homologs are encoded in prokaryote genomes, and up to five in non-photosynthetic eukaryotes, the have proliferated particularly in plants. Seventeen homologs are encoded in the model plant, *Arabidopsis thaliana*, six of which localize to chloroplast plastoglobules where ubiquinone is neither biosynthesized nor accumulated. This suggests that these proteins have roles related to metabolism of other quinone and prenyl lipid compounds, likely including multiple compounds employed in photosynthesis (7).

In spite of the significance of the ABC1 family to organismal survival across a wide evolutionary space, it is still disputed whether they are bona fide protein kinases. With limited sequence conservation to the “classical” eukaryotic protein kinases (ePKs), the ABC1s nevertheless have several conserved key kinase motifs, such as the DFG motif responsible for divalent cation coordination and nucleotide binding, and the invariant Lys that binds and transfers the γ- phosphate. However a number of other key motifs are missing, and multiple conserved motifs unique to ABC1s, with unclear catalytic roles, are evident (7, 10). Additionally, the Gly-rich loop, central in ATP binding, is found instead as a conserved Ala triad (AAAS sequence) in what is known as the A-rich loop.

It has been difficult to demonstrate conclusive protein kinase activity among ABC1 proteins. Protein kinase activity may be facilitated in ABC1s through unique catalytic or regulatory mechanisms that demand unique conditions. Alternatively, protein kinase activity may be precluded by the unique features of the ABC1 domain, existing instead as a so-called unorthodox kinase, i.e. pseudokinase or lipid/metabolite kinase. On the one hand, the crystal structure of a truncated sequence of the human ABC1 ortholog, HsADCK3^NΔ254^, revealed a classical bi-lobed protein kinase architecture but with a KxGQ domain, unique to the ABC1 family, laying over the active site and preventing access of substrate (11) (**SI Appendix, Fig. S1**). No protein kinase activity was found with HsADCK3^NΔ254^ by *in vitro* assay, but curiously, an A-to-G point mutant in the Ala triad (A339G) did exhibit clear protein kinase activity (11). In addition, phosphoproteomics experiments with mice either did not identify evidence of phosphorylation on ubiquinone biosynthesis proteins, or did not find a statistically significant decrease of phosphorylation in the absence of the ABC1 homolog (12). Collectively, the authors concluded that endogenous HsADCK3 represents a pseudokinase capable of ATPase consumption but not protein kinase activity (11–13). On the other hand, *in vitro* assays with the plastoglobule-localized AtABC1K1 and AtABC1K3 provided evidence for weak trans-phosphorylation of the plastoglobule-localized substrate, tocopherol cyclase (14, 15). Our own recent work has also demonstrated indirect evidence for *in vitro* ABC1 kinase activity associated with isolated *Arabidopsis thaliana* plastoglobules that is substantially reduced in an *abc1k1/abc1k3* double mutant (5). Furthermore, isoelectric point shifts on several ubiquinone biosynthesis proteins in yeast, ascribed to phosphorylation, appeared in a COQ8/ABC1-dependent manner (8). These shifts could also be rescued by heterologous expression of HsADCK3. These results support the idea that plastoglobule-localized and other ABC1 proteins hold protein kinase activity. Here, we report our investigation of the plastoglobule-localized AtABC1K6. We demonstrate that with proper buffer conditions clear protein kinase activity is exhibited as autophosphorylation and that this activity is associated with a functional role in plant development. These results provide strong support for a central regulatory role of the ABC1s on plastoglobules and present the experimental means to investigate the targets, functions, and regulation of ABC1s.

## RESULTS

### Loss of AtABC1K6 impairs the transition from vegetative to reproductive growth

The *A. thaliana* plastoglobule harbors six members of the ABC1 protein family. A co-expression network of plastoglobule genes previously demonstrated that AtABC1K6 was positioned as a network hub, suggesting a central role in the regulation of the plastoglobule (6). To investigate the function of AtABC1K6 we isolated a SALK T-DNA insertion mutant line in the sixth exon of the AtABC1K6 gene (At3g24190, hereafter referred to as *abc1k6-1)*, which fails to accumulate detectable transcript (**SI Appendix, Fig. S2**). Analysis of photosynthetic activity did not reveal a difference between wild-type (Columbia-0) and *abc1k6-1* (**SI Appendix, Fig. S3**). However, under standard growth chamber conditions, we observed that *abc1k6-1* plants bolted and flowered several days later than the wild-type plants (**Fig. 1A**). The developmental rates of *abc1k6-1* and wild-type were quantified according to the *A. thaliana* developmental stage descriptors established by Boyes et al. (16), which demonstrated that *abc1k6-1* reached stage 5.10 and stage 6.0 (first flower buds and first flower opening, respectively) around three days later than wild-type, whether grown under long day (LD; 16/8 hours photoperiod) or constant light (CL) conditions (**Fig. 1B**). Furthermore, under CL, mutant plants were also delayed in reaching stage 6.9 (end of senescence) compared to the wild-type. In contrast, the rate of leaf emergence and the rosette size throughout vegetative growth was indistinguishable. Furthermore, the time from bolting (stage 5.10) to first flower open (stage 6.0) was not altered in *abc1k6-1*. Together, this indicates that *abc1k6-1* is impacted specifically in the developmental transition from vegetative to reproductive growth, rather than the rate of vegetative or reproductive development per se.

**Fig. 1.**
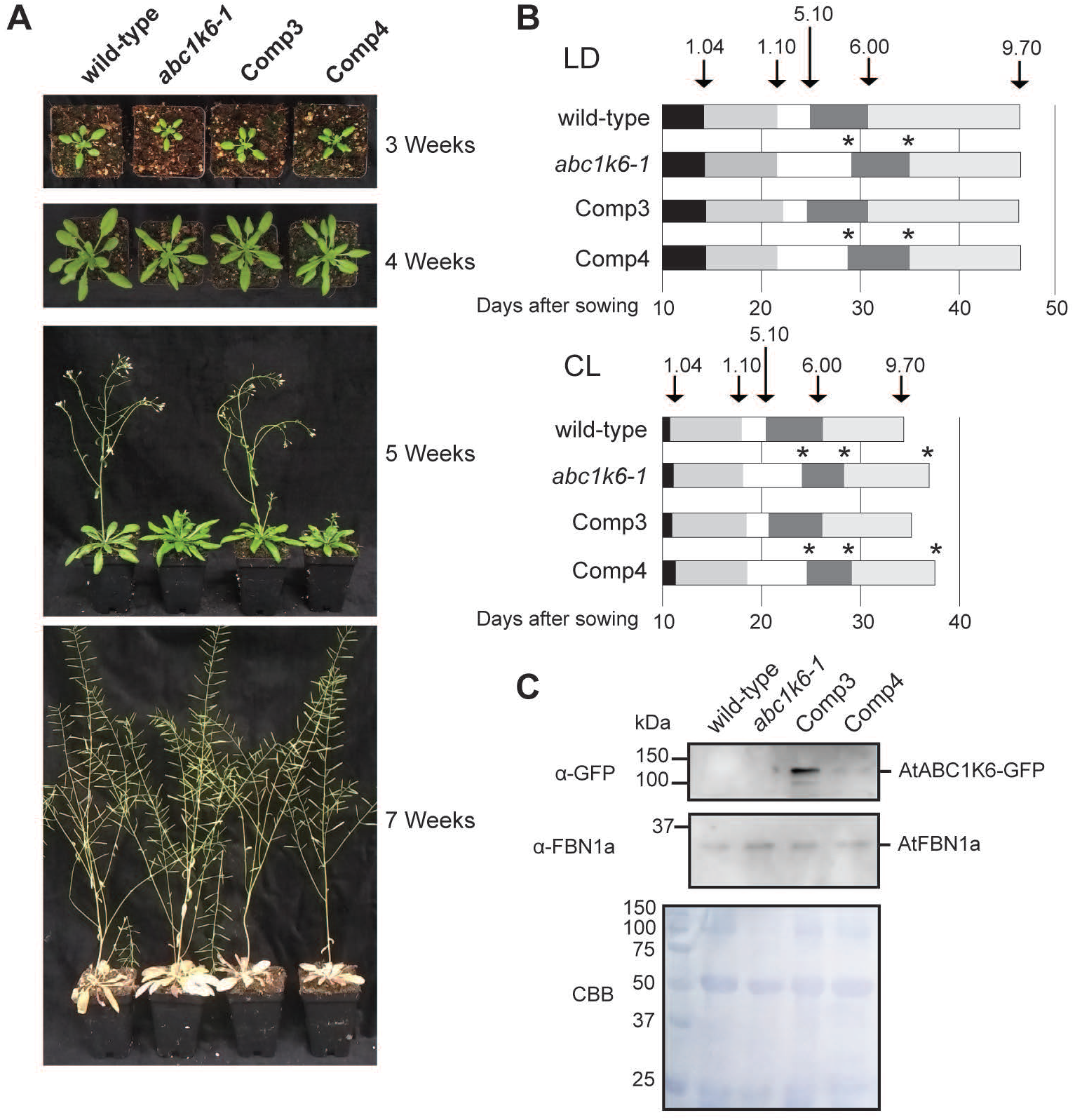
The *abc1k6-1* T-DNA lesion impairs proper initiation of reproductive growth. (A) *abc1k6-1* is delayed in bolting and flowering. Plants were grown on soil in a growth chamber with LED lighting under long day (LD: 16 hours light and 8 hours dark, 120 μmol m^-2^ s^-1^) or constant light (CL; 120 μmol m^-2^ s^-1^) photoperiod. Representative plants grown under LD photoperiod are shown at selected time points during development. (B) Schematic illustrating the developmental stage transitions of each genotype following the nomenclature described by Boyes et al., 2001 (17), given as the average time from 10 individual plants. Arrows indicate the time in which wild-type plants reached the respective developmental stage: 1.04: four rosette leaves; 1.10: 10 rosette leaves; 5.10: first flower buds visible; 6.00: first flower open; 9.70: senescence complete. Asterisks indicate a statistically significant difference (paired, two-tailed Student’s T-test with p-value less than 0.05; n = 10) in reaching a given developmental stage relative to wild-type. (C) Immunoblot of total protein extracts from *A. thaliana* leaves. Anti-GFP antibody is used to quantify expression of the AtABC1K6-GFP construct. Rabbit anti-FBN1a/PGL35 antibody is used as plastoglobule marker protein to estimate the prevalence of plastoglobules in each line. Coomassie brilliant blue (CBB) stain of the same membrane is provided as a loading control.

To verify that this developmental phenotype is associated with disruption of AtABC1K6, we transformed *abc1k6-1* with AtABC1K6 cDNA fused C-terminally to GFP under the cauliflower mosaic virus 35S promoter (**Fig. 1C**). Introduction of AtABC1K6-GFP fully complemented the developmental phenotype as seen in the Complementation line 3 (Comp3) which over-expressed the ABC1K6-GFP product (**Fig. 1**), confirming that the delayed developmental transition arises due to loss of AtABC1K6. In investigating the sub-chloroplast localization of the AtABC1K6-GFP fusion in complementation line 3 (Comp3) we discovered that the fusion protein was predominantly associated with thylakoid and stromal fractions when assaying with anti-GFP antibody (**SI Appendix, Fig S4**). Similarly, confocal microscopy results showed a diffuse localization of GFP fluorescence throughout chloroplasts. However, our proteomic analyses of isolated plastoglobules from wild-type, *abc1k6-1* and Comp3 (see below) revealed that AtABC1K6-GFP is also prevalent in plastoglobule samples in Comp3, and that its magnitude of overaccumulation in leaf tissue matches its overaccumulation in plastoglobule samples (**Fig. 2A**). We speculate that the high expression level of AtABC1K6-GFP causes spill-over of the heterologous protein from its natural location at the plastoglobule leading to ectopic accumulation in additional sub-compartments of the chloroplast.

**Fig 2.**
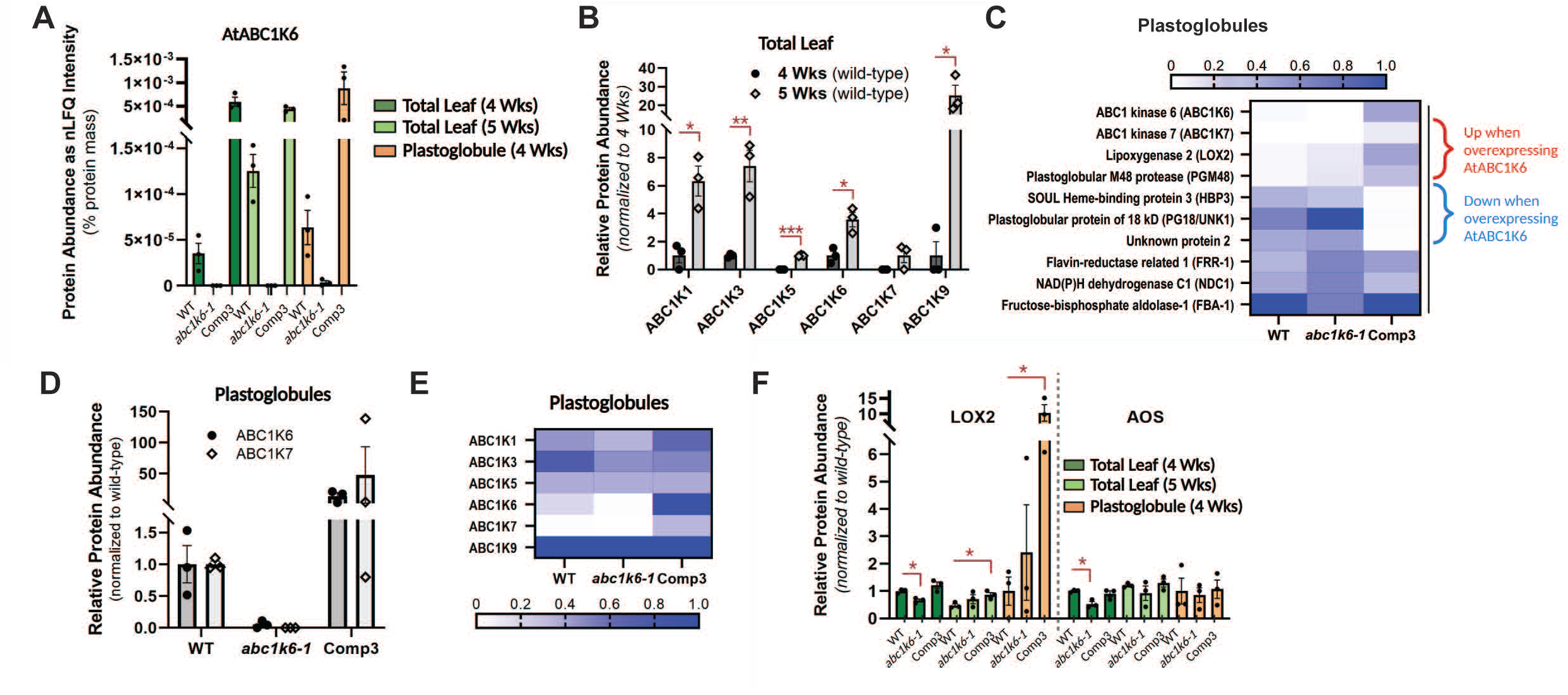
Alterations to the leaf and plastoglobule proteomes of *abc1k6-1* and Comp3. (A) The relative abundance of AtABC1K6 in leaf and plastoglobule samples as determined by label-free quantitative (LFQ) proteomics, given as normalized LFQ (nLFQ). The fold-change from wild-type to Comp3 is about the same in leaf tissue and plastoglobules at 4 weeks. (B) The six plastoglobule-localized ABC1K homologs increase in wild-type leaf tissue from week 4 to week 5. Each protein’s abundance is normalized to the week 4 level, except for AtABC1K5 and AtABC1K7 which are not detected at week 4. (C) A heatmap of plastoglobule proteins impacted by loss of AtABC1K6 illustrating the changes in the plastoglobule proteome at week 4. All plastoglobule proteins that are identified as being impacted by loss of AtABC1K6 are presented. Values are normalized to the highest value among the dataset (i.e. FBA-1 in wild-type). (D) Levels of AtABC1K6 and AtABC1K7 in the plastoglobule proteome closely track with each other across each of the genotypes. Each protein is normalized to its wild-type value. (E) A heatmap illustrates the changes of each plastoglobule-localized ABC1 homolog in the plastoglobule proteome at week 4. Values are normalized to the highest value among the dataset (i.e. ABC1K9 in wild-type). (F) Protein quantification suggests Lipoxygenase 2 of the JA biosynthetic pathway is remobilized to the plastoglobule in *abc1k6-1*. Two plastoglobule-localized enzymes of jasmonic acid biosynthesis were reduced in leaf tissue of *abc1k6-1* at week 4, while the lipoxygenase specifically was elevated about 10-fold at the plastoglobule in the presence of over-expressed AtABC1K6 in the Comp3 line. Values are normalized to each protein’s wild-type week 4 level (total leaf samples) or to the wild-type level (plastoglobule samples). In all panels, n = 3 biological replicates; error bars indicate ±1 standard error of the mean; where error bars are not shown they are smaller than the data point. In panels, B and F, significant differences between samples are indicated with a red line and/or asterisks; one asterisk denotes p-value <0.05, two asterisk denotes p-value < 0.01, three asterisks denote a p-value < 0.001.

### Plastoglobule-localized ABC1 proteins are induced during development

To test how changes to the leaf proteome may influence the developmental phenotype we performed quantitative proteomic analyses of leaf samples from wild-type, *abc1k6-1*, and Comp3, at 4 weeks and 5 weeks (28 and 35 days after germination [DAG], respectively) in LD conditions. These time points represent several days before and several days after wild-type has initiated bolting (stage 5.10), providing an easily discernible developmental marker of the transition from vegetative to reproductive growth. The quantitative proteomic analysis identified and quantified 4076 proteins or protein groups (hereafter referred to collectively as proteins) across all samples (**SI Appendix, Dataset S1**). Quantification of proteins employed label-free quantification (LFQ) of MS1 peptide ion intensity as implemented in the MaxQuant proteomics software pipeline (17, 18). To determine those proteins whose abundance was effected by the mutation, we identified proteins that fit both of the following criteria: i) a significantly different normalized LFQ (nLFQ) in *abc1k6-1* relative to wild-type, [p-value <0.05, controlling for multiple hypotheses using the Benjamini-Hochberg method (19)] and ii) an nLFQ restored at least half-way to wild-type levels in Comp3. With this in mind, a total of 313 proteins were affected at 4 weeks and 379 proteins at 5 weeks. Surprisingly, only 56 of these proteins were affected at both time points, indicating effects on the proteome are highly dependent on developmental stage.

As expected, AtABC1K6 could be readily identified and quantified in wild-type and Comp3 genotypes, but not in *abc1k6-1* (**SI Appendix, Dataset S1**). From the proteomics results we could estimate that AtABC1K6 accumulation in leaf tissue was about 16-fold higher than wild-type levels at 4 weeks, and about 3-fold higher at 5 weeks (**Fig. 2A**). The greater differential at 4 weeks was due to a tripling of endogenous AtABC1K6 levels in wild-type from week 4 to week 5. Interestingly, we found that the other plastoglobule-localized ABC1 homologs also were significantly increased from week 4 to week 5, in particular AtABC1K9 whose accumulation increased over 25-fold (**Fig. 2B**). We also looked for the accumulation patterns of proteins with well-known roles in flowering (e.g. Flowering Locus T, Flowering Locus Y, Suppressor of Overexpression of Constans 1, and Leafy). However, these proteins were not identified in our dataset, presumably because of low abundance within the leaf proteomes that prevented their detection with the LC-MS/MS.

Significantly, three of the enzymatic steps of jasmonic acid (JA) biosynthesis were significantly reduced in *abc1k6-1* at week 4, but not week 5 (**Table 1**). JA signaling can promote chlorophyll degradation and senescence, and the plastid-localized steps have been found to associate with plastoglobules under specific conditions (4, 20–26). We considered whether a possible deficiency in JA synthesis, and hence signaling, at week 4 may account for the developmental phenotype in *abc1k6-1*. To test this possibility, we treated wild-type and *abc1k6-1* plants with 25 μM of Me-JA as a foliar spray occurring each day for seven days from 21 to 28 DAG. While leaves of the treated samples accumulated anthocyanins, indicating perception of JA, the time at which each genotype reached bolting was unchanged relative to mock-treated plants, indicating the JA treatment failed to complement the developmental phenotype (**SI Appendix, Fig S5**).

**Table 1.** Three Proteins of Jasmonic Acid Biosynthesis Are Depleted in Leaf Tissue at Week 4 in *abc1k6-1*

|  |  |  |  |
| --- | --- | --- | --- |
| **Accession** | **Annotation** | **Location** | **Fold-change<sup>a</sup> ($\log_2$ of *k6*/WT)** |
| AT3G45140.1 | Lipoxygenase (AtLOX2) | Plastid - Plastoglobules; Envelope | -0.60 |
| AT5G42650.1 | Allene oxide synthase (AtAOS) | Plastid - Plastoglobules; Envelope | -0.92 |
| AT2G06050.1 | 12-Oxophytodienoate Reductase 3 (AtOPR3) | Peroxisome | -2.15 |

<sup>a</sup> the three proteins depleted in *abc1k6-1* are all complemented (at least half restored to wild-type levels) in the *Comp3* line, indicating that their change in abundance is due to loss of *AtABC1K6*.

### Quantitative proteomics reveals specific changes to the plastoglobule proteome

Comparison of protein localizations among the total leaf proteomes revealed that the sum of plastoglobule protein levels were reduced by about 20% relative to wild-type, which cannot be accounted for solely by the loss of AtABC1K6 (**SI Appendix, Dataset S2**). To identify specific changes to the plastoglobule core proteome associated with loss of AtABC1K6 we compared the proteomes of isolated plastoglobules from wild-type, *abc1k6-1* and Comp3 at week 4 (6). Ten proteins of the plastoglobule core proteome were effected by loss of AtABC1K6, according to the same criteria used with leaf samples (**Fig. 2C, Dataset S3**). Only a single tryptic peptide of AtABC1K6 was identified in *abc1k6-1* plastoglobules, which came from near the N-terminus (residues 175-183), upstream of the T-DNA insertion site. This suggests truncated AtABC1K6 accumulates at low levels and associates with the plastoglobule in *abc1k6-1*. About 14-fold more protein was associated with plastoglobules in Comp3 than wild-type, consistent with the enrichment of AtABC1K6 seen at the leaf tissue level (**Fig. 2A**). Strikingly, nLFQ levels of AtABC1K7, but not the other plastoglobule-localized ABC1 homologs, closely tracked with that of AtABC1K6 over the genotypes; no AtABC1K7 was found in isolated plastoglobules of *abc1k6-1*, while AtABC1K7 associated with plastoglobules to nearly 50-fold higher levels in Comp3 than in wild-type (**Fig. 2 D and E**). This suggests that AtABC1K6 may stabilize AtABC1K7, or be necessary for its association with plastoglobules. Lipoxygenase 2 (AtLOX2), involved in JA biosynthesis, and Plastoglobular Metalloprotease M48 (AtPGM48), a positive regulator of leaf senescence, also were more abundant in plastoglobule samples of Comp3 (**Figure 2C**). The 10-fold enrichment of AtLOX2 at the Comp3 plastoglobule was not reflected at the leaf level where total levels of AtLOX2 were the same at the week 4 time point when plastoglobules were isolated (**Fig. 2F**). This is indicative of a recruitment of AtLOX2 to the plastoglobule upon overaccumulation of AtABC1K6-GFP, reminiscent of recruitment of this and other JA biosynthesis enzymes to the plastoglobule in other conditions (4, 5). In contrast, evidence did not support recruitment of AtAOS and AtPGM48 to the plastoglobule, since AtAOS levels on the plastoglobule were comparable in all genotypes, while AtPGM48 levels were below the detection limit in leaf samples (**Fig. 2F**). Curiously, three proteins were conspicuously absent from plastoglobules of Comp3, despite their prominent accumulation in plastoglobule samples of wild-type and *abc1k6-1*: i) a SOUL heme-binding protein (AtHBP3), ii) the Plastoglobule protein of 18 kDa (AtPG18), and iii) an uncharacterized protein called Unknown 2 (**Fig. 2C**). The elevated levels of AtPGM48 metalloprotease, in tandem with the specific loss of the above three proteins, presents these three proteins as candidate substrates for proteolytic cleavage by AtPGM48. None of the three proteins have known functions, although it seems unlikely that their absence from plastoglobules is related to the phenotypic complementation, since they are prominent components of the wild-type plastoglobules.

### AtABC1K6 is a protein kinase that requires low salt and Mn^2+^ as co-factor

Multiple conserved residues and kinase motifs shared with canonical ePKs suggest that the ABC1 family encodes protein or lipid kinase activity, however clear demonstration of any kinase activity remains elusive (11, 13). To test for protein kinase activity of AtABC1K6 we heterologously expressed the mature AtABC1K6 protein (i.e. lacking only the chloroplast transit peptide; residues 1-19) in *Escherichia coli*, fused at the N-terminus to the maltose-binding protein (MBP), containing a Factor Xa protease cleavage site, and at the C-terminus to a 6xHis tag (MBP-AtABC1K6-His; **Fig. 3C**). Purified MBP-AtABC1K6-His was incubated in *in vitro* kinase buffer with γ-^32^P-labelled ATP for 30 minutes and separated with SDS-PAGE. No *in vitro* kinase activity could be detected in the presence of 150 mM NaCl (data not shown). However, in buffer lacking NaCl, *in vitro* kinase activity was readily apparent from a spontaneous cleavage product of the recombinant protein (**Fig. 3A**). This cleavage product appeared during the purification despite our exclusion of Factor Xa or any other protease. Mass spectrometry-based proteomic analysis of the cleaved protein fragment, separated by SDS-PAGE, identified the cleavage site after K443 (counting from the 1^st^ residue of the MBP-AtABC1K6-His recombinant protein) (**Fig. 3C**). This indicates the cleavage product receiving γ-^32^P contains AtABC1K6 protein minus the 71 N-terminal residues which eliminates the a predicted intrinsically disordered region (**SI Appendix, Fig. S6**). This conclusion is consistent with the immunoblotting data that indicates that the cleavage product retains the 6xHis tag but not the MBP tag, and its migration at ca. 75 kD is consistent with the theoretical molecular weight of the putative protein fragment, calculated at 80.7 kD. Hence, we refer to this kinase-active cleavage product as AtABC1K6^NΔ71^-His for the remainder of this article. Importantly, full-length MBP-AtABC1K6-His migrating at ca. 130 kD did not display kinase activity, suggesting that either the presence of the MBP tag or the disordered region inhibits the kinase activity in the *in vitro* system.

**Fig. 3.**
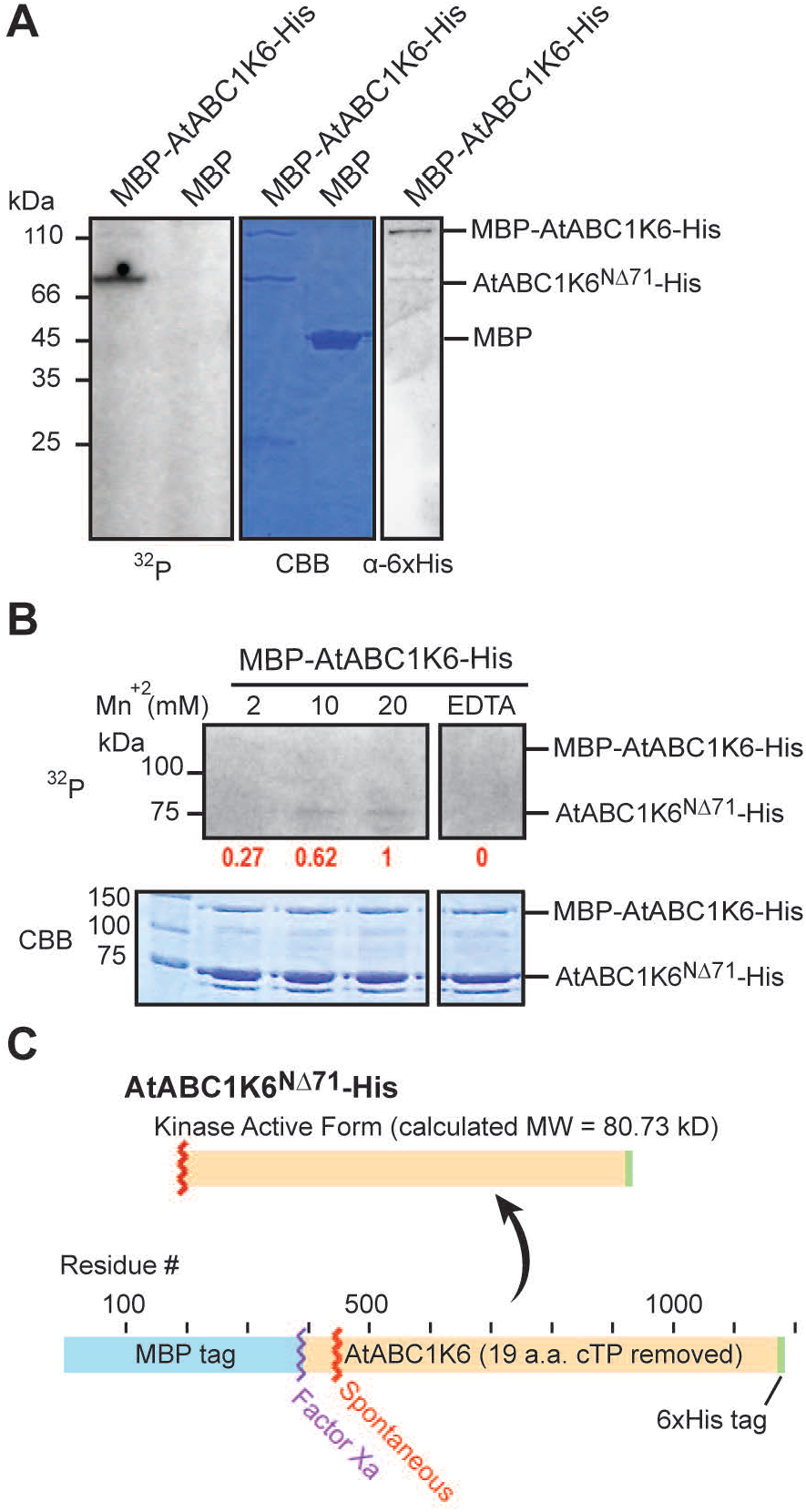
AtABC1K6 displays *in vitro* kinase activity dependent on Mn^2+^. (A) AtABC1K6 protein fused to MBP was expressed in *E. coli* and purified to perform *in vitro* autophosphorylation assays using γ-P^32^-ATP. On the left panel is shown the autophosphorylation activity of the heterologously expressed protein, while MBP alone was used as a negative control. A spontaneously occurring, truncated form of MBP-AtABC1K6-His migrates at ca. 75-80 kD, is indicated as ‘AtABC1K6^NΔ71^-His’, and displays *in vitro* kinase activity. Two μg of protein was loaded in each lane. Samples were incubated with γ-P^32^-ATP for 30 minutes and gel was exposed to autoradiography film for 7 days. The Coommassie stained membrane (CBB) is indicated as a loading control. (B) The purified MBP-AtABC1K6-His was assayed for in-vitro kinase activity with increasing concentrations of Mn^2+^ in buffer containing 20 mM Tris (pH 7.5) and no NaCl. As a negative control for kinase activity, EDTA was added to the assay with 20 mM Mn^2+^. Sample was incubated with γ-P^32^-ATP for 45 minutes and the gel was exposed to autoradiography film for 3 days. The two spliced panels are from the same image of the same film/gel. The values in red indicate the relative intensity of radiography at the AtABC1K6^NΔ71^-His band, quantified using Image J, and normalized to the 20 mM Mn^2+^ lane. Five μg of protein was loaded in each lane. (C) The identity of the spontaneous cleavage product, and location of the spontaneous cleavage, was made by excising bands from an SDS-PAGE gel and analyzing by LC-MS/MS, as described in the main text and illustrated in **Fig. S6**. A schematic illustration of the MBP-AtABC1K6-His recombinant protein, and the locations of the encoded Factor Xa cleavage site and the spontaneous cleavage site occurring during expression or purification is given.

To closely mimic the *in vivo* stromal environment of AtABC1K6, we initially used 5 mM Mg^2+^ and 0.5 mM Mn^2+^ in our *in vitro* kinase buffer, concentrations matching those estimated in chloroplast stroma during photosynthesis (27–29). We then tested whether the kinase activity depended on the Mg^2+^ or Mn^2+^ by providing only one of the two cations. Assays revealed dose-dependent activity in the presence of Mn^2+^, however no kinase activity could be detected even at the highest tested concentration of Mg^2+^ (**Fig. 3B, SI Appendix Fig. S7**). We conclude that a truncated form of AtABC1K6, lacking a predicted disordered region at its N-terminus, encodes protein kinase activity that employs Mn^2+^ as co-factor and is regulated by ionic strength of the buffer.

### Kinase activity is associated with isolated plastoglobules

We have previously demonstrated *in vitro* protein kinase activity with isolated plastoglobules that target multiple substrate proteins of varying molecular weights (5). In order to understand the cation dependency in the context of purified plastoglobules, we tested *in vitro* kinase activity with wild-type plastoglobules under different cation concentrations. Interestingly, although 20 mM Mn^+2^ showed detectable kinase activity, a stronger signal was obtained under 20 mM of Mg^+2^, while no kinase activity was observed under 20 mM of Ca^+2^ (**Fig. 4A**). We then performed *in-vitro* kinase assays using isolated wild-type plastoglobules in kinase buffer with gradients of NaCl and KCl. Consistent with results from purified AtABC1K6, kinase activity associated with plastoglobules was sensitive to both salts, preferring low salt concentrations of less than 50 mM (**Fig. 4B**). To determine if any of the detectable protein kinase activity from isolated plastoglobules might derive from AtABC1K6, we compared protein kinase activity of plastoglobules and chloroplasts isolated from wild-type and *abc1k6-1*. However, band patterns were essentially unchanged between the two genotypes, indicating AtABC1K6 does not account for the detectable protein kinase activity in our *in vitro* kinase assays with plastoglobules (**Fig. 4C**). Thus, the salt sensitivity appears to be a general phenomenon among plastoglobule-localized ABC1 paralogs, while the requirement for Mn^2+^ as divalent cation co-factor is not absolute across all plastoglobule-localized ABC1 homologs.

**Fig. 4.**
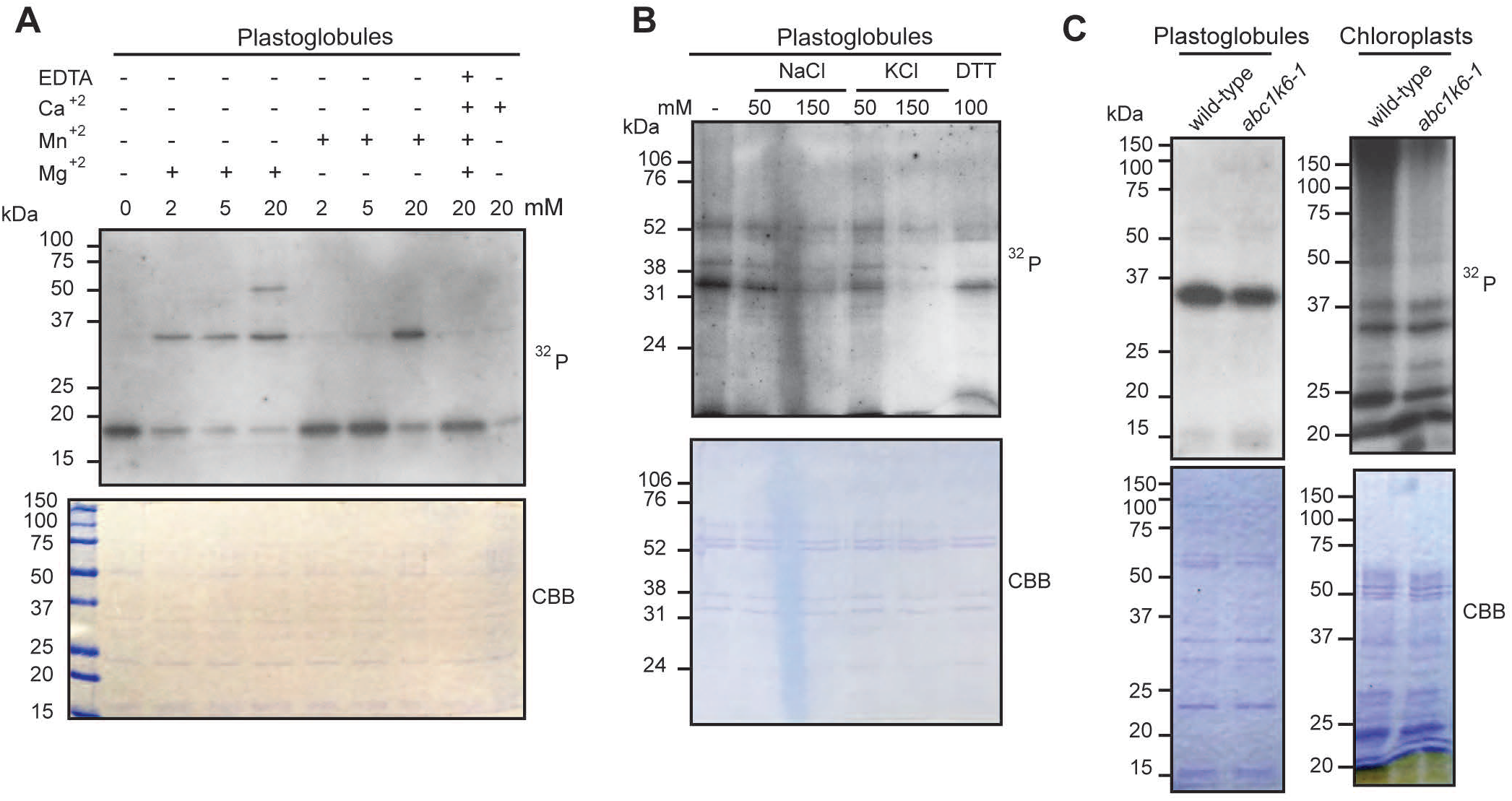
Isolated plastoglobules demonstrate *in vitro* kinase activity that is dependent on salt concentration. (A) Kinase activity associated with plastoglobules of wild-type *A. thaliana* can use Mn^2+^ or Mg^2+^ as co-factor. Detectable activity primarily is seen at ca. 35 kD, which may be one or more of the prominent FBN proteins associated with plastoglobules. Sample was incubated with γ-P^32^-ATP for 45 minutes, and gel was exposed to autoradiography film for 24 hours. (B) Plastoglobules of wild-type *A. thaliana* are sensitive to higher salt concentrations, as seen with the AtABC1K6 recombinant protein, but not to reducing environment. Sample was incubated with γ-P^32^-ATP for 45 minutes, and gel was exposed to autoradiography film for 48 hours. (C) The kinase activity of isolated plastoglobules and whole chloroplasts are not impacted by the loss of AtABC1K6, as revealed in the activity assays with the *abc1k6-1* line. Sample was incubated with γ-P^32^-ATP for 45 minutes, and gel was exposed to autoradiography film for 24 hours. Assays in (A) and (C) were performed in buffer lacking NaCl and KCl.

### *In vitro* kinase activity and *in vivo* functionality of AtABC1K6 is dependent on the conserved Ala triad of the ATP-binding pocket

A unique feature of the ABC1 protein kinase domain is the substitution of a highly conserved Ala triad in the ATP-binding pocket, in place of the canonical GxGxxG motif (the Gly loop, where x indicates any residue) seen in ePKs (**Fig. 5B**) (7, 10, 30). We evaluated the role of the Ala triad in AtABC1K6 with *in vitro* kinase assays using point mutants in the triad. As with the wild-type recombinant protein, point mutants were spontaneously cleaved during purification and migrated at the same apparent molecular weight. Strikingly, the A268G mutation (mutated in the first Ala of the triad, equivalent to the A339G mutation of HsADCK3^NΔ254^), showed severely diminished kinase activity (**Fig. 5A**). Furthermore, point mutants of the next two Ala residues (A269G and A270G) and the triple A-G point mutant also showed severely diminished kinase activity.

**Fig. 5.**
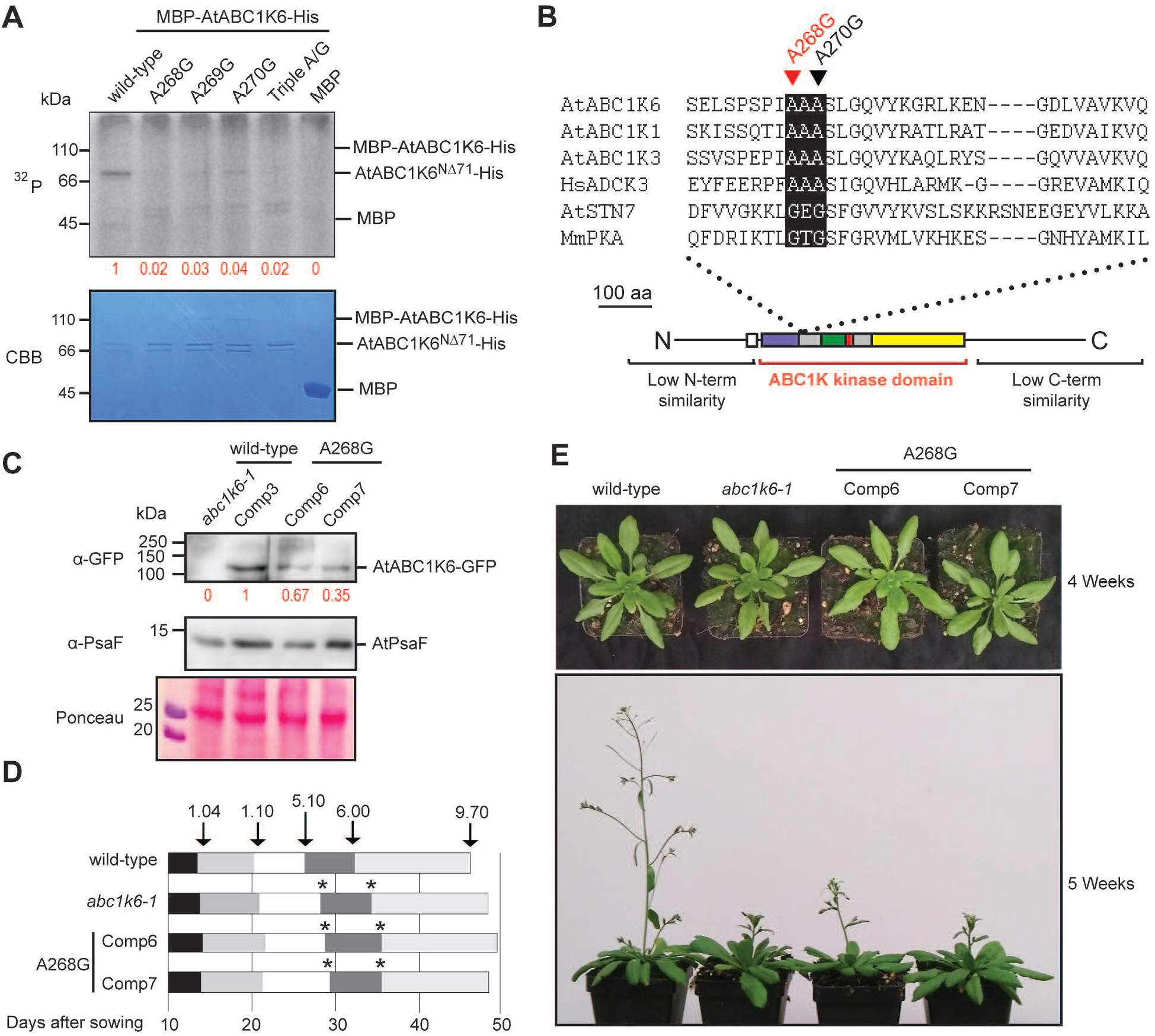
The highly conserved Ala triad of the ABC1K kinase domain is essential for *in vitro* kinase activity and *in vivo* function of AtABC1K6. (A) *In vitro* kinase assays with the wild-type sequence, and various point mutants of the Ala triad, of MBP-AtABC1K6-His are presented. The values in red indicate the relative intensity of radiography at the AtABC1K6^NΔ71^-His band, quantified using Image J, and normalized to the wild-type lane. Three μg of protein was loaded in each lane. Sample was incubated for 30 minutes with γ-P^32^-ATP and the gel was exposed to autoradiography film for 1 week. (B) A sequence alignment of several ABC1 homologs, along with two canonical protein kinases, AtSTN7 and MmPKA from *A. thaliana* and *Mus musculus*, respectively. The alignment is intended to illustrate the sequence context of the point mutants tested in (A) and illustrate the conservation of the unique ATP-binding pocket of the ABC1 family. AtABC1K6, AtABC1K1, and AtABC1K3 are plastoglobule-localized ABC1 homologs from *A. thaliana*, HsADCK3 is an ABC1 homolog from mitochondria of *Homo sapiens*, AtSTN7 is State Transition 7 kinase of *A. thaliana* thylakoid membranes, and MmPKA is Protein Kinase Alpha of *M. musculus*. (C) An anti-GFP immunoblot demonstrates expression of the AtABC1K6-GFP in various lines stably transformed with either the wild-type (Comp3), or A268G point mutant (Comp6 & Comp7) variant of AtABC1K6. Anti-PsaF, detecting a membrane protein of the photosystem II complex, was used as a loading control. Values in red represent the relative ratio of GFP intensity normalized by PsaF. (D) Transformation with the point mutants fails to complement the developmental phenotype, in contrast to transformation with the wild-type sequence. Schematic illustrates the developmental stage transitions of each genotype following the nomenclature described by Boyes et al., 2001 (16), given as the average time from 10 individual plants. Arrows indicate the time in which wild-type plants reached the respective developmental stage: 1.04: four rosette leaves; 1.10: 10 rosette leaves; 5.10: first flower buds visible; 6.00: first flower open; 9.70: senescence complete. Asterisks indicate a statistically significant difference in reaching a given developmental stage compared to wild-type. n = 10; Student’s T-test with p-value less than 0.05. (E) Representative photos of genotypes demonstrating the lack of complementation seen in the lines transformed with the A268G point mutant.

To test if the *in vitro* kinase activity of AtABC1K6 is relevant to *in vivo* function we attempted complementation of *abc1k6-1* with the A268G point mutant of AtABC1K6. To mimic the conditions of the Comp3 line that fully complemented the developmental phenotype, we retained the C-terminal GFP tag and 35S promoter sequences. Despite strong overexpression of the A268G point mutant in the *abc1k6-1* background, no complementation of the developmental phenotype was apparent (**Fig. 5C, D, and E**), contrasting with the full complementation seen when using the wild-type sequence in Comp3 (**Fig. 1**). This indicates the *in vitro* kinase activity detected with AtABC1K6^NΔ71^-His recombinant protein is necessary for its *in vivo* function related to the developmental transition.

### The human ABC1 ortholog, HsADCK3^NΔ254^, also exhibits protein kinase activity but with differing requirements

In light of the distinct requirements for protein kinase activity that we found with AtABC1K6, we considered whether previous efforts may not have detected kinase activity with HsADCK3^NΔ254^ due to the chosen buffer conditions. Thus, we tested HsADCK3^NΔ254^ for divalent cation dependency and salt sensitivity. As for AtABC1K6, HsADCK3^NΔ254^ exhibited protein kinase activity when Mn^+2^ was provided as divalent cation (**Fig. 6A**). However, activity was also clearly seen when providing solely Mg^+2^, albeit not as strong as with Mn^2+^. This indicates that HsADCK3^NΔ254^ is more active with Mn^+2^ but can also accommodate Mg^2+^ as divalent cation co-factor, contrasting with the apparent absolute requirement for Mn^2+^ seen with AtABC1K6.

**Fig. 6.**
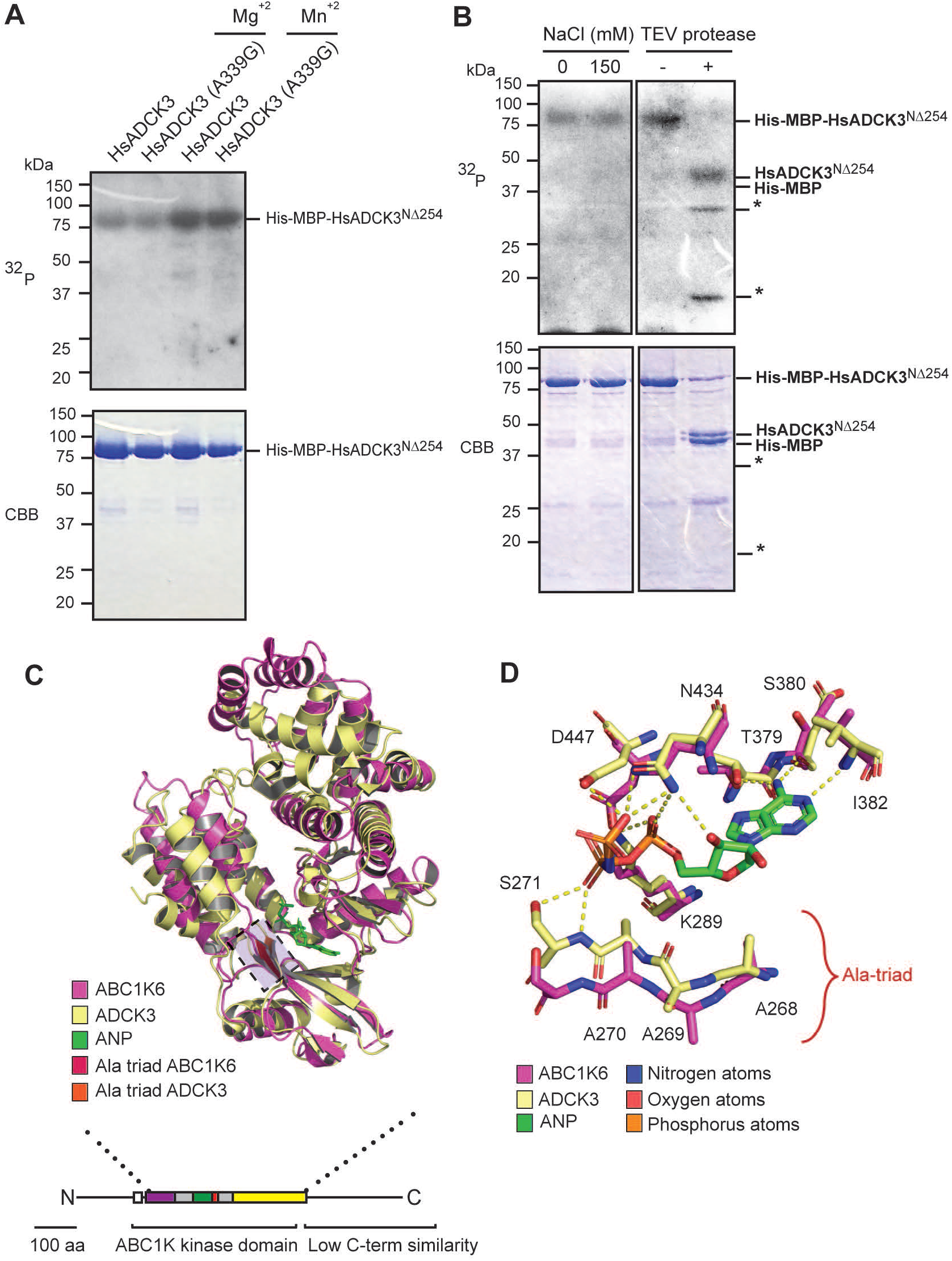
HsADCK3^NΔ254^ is capable of autophosphorylation in *in vitro* kinase assays with Mn^2+^ or Mg^2+^, and independent of salt concentration. (A) The wild-type and point mutant variant of HsADCK3^NΔ254^ was capable of autophosphorylation in the presence of either divalent cation. Purified protein was incubated with γ-P^32^-ATP, 20 mM Tris-HCl (pH 7.5), no NaCl, and either 20 mM Mn^2+^ or Mg^2+^ for 45 minutes and exposed to autoradiography film for 3 days. Five μg of protein was loaded in each lane. Coomassie staining (CBB) of an independent gel is used as loading control. (B) HsADCK3^NΔ254^ is insensitive to salt, displaying equally strong *in vitro* kinase activity at 0 and 150 mM NaCl. In the left panel, full length recombinant protein was assayed in the presence or absence of NaCl. In the right panel the protein was assayed in the absence of salt using the full length protein and cleaved product using the TEV protease cleavage site downstream of the His-MBP tag. Samples were incubated with γ-P^32^-ATP for 45 minutes and gel was exposed to autoradiography film for 24 hours. Three μg of protein was loaded in each lane. Coomassie staining (CBB) of the same membrane is used as loading control. (C) Alignment of the ABC1 domains of HsADCK3 (PDB 5i35, yellow) and AtABC1K6 (Alphafold AF-Q9LRN0-F1, magenta) reveals strong structural conservation. The Ala-triad and β-strand are highlighted with a lavender colored box. The RMSD of the alignment is 2.868. (D) Close-up view of the Ala-triad and ATP binding pocket from the alignment shown in panel C. Residue numbering follows the AtABC1K6 full-length protein sequence.

When testing salt sensitivity in the presence of both 5 mM Mg^2+^ and 0.5 mM Mn^2+^ the recombinant HsADCK3^NΔ254^ surprisingly displayed clear *in vitro* kinase activity at both 0 and 150 mM NaCl, whether the MBP tag is retained or cleaved off (**Fig. 6B**). This contrasts with previous results from Stefely, et al. that did not identify *in vitro* kinase activity with HsADCK3^NΔ254^ at 150 mM NaCl and 20 mM Mg^2+^ (11, 13).

Both the wild-type and A339G point mutant of HsADCK3^NΔ254^ displayed strong kinase activity, indicating the HsADCK3^NΔ254^ ortholog does not share the delicate sensitivity of the Ala triad seen in the AtABC1K6 under our *in vitro* kinase conditions (**Fig. 6A**). These results indicate possible structural differences in the ATP-binding pocket between the *A. thaliana* and human orthologs that may also account for the differential sensitivity to salt concentration shown above. To investigate this possibility, we compared the crystal structure of HsADCK3^NΔ254^ (PDB: 5i35) with a modeled structure of AtABC1K6 generated using Alphafold (ID: AF-Q9LRN0-F1) which generates models de novo, i.e. independent of any homology template (31). Three-dimensional alignment of the kinase domain structures from AtABC1K6 and HsADCK3 revealed considerable architectural conservation (Root-mean-square deviation of 2.868 for atomic positions 1305 to 1305 atoms) (32). Both structures revealed that the Ala-triad forms the last strand of a β-sheet which cups the ATP analog (**Fig. 6C**). The similar structural features of the ATP-binding pocket preclude an explanation for the differing effects of the A-to-G mutations. However, the adjacent conserved Ser is positioned to form a hydrogen bond with the β-phosphate of the ATP (**Fig. 6D**). Hence, the β-sheet, comprised of the Ala-triad, is important for the positioning of the Ser residue and stability of the ATP. We suggest that the provision of Gly within this triad introduces an enhanced flexibility which destabilizes the strand and removes a part of the ATP-binding pocket explaining the loss of kinase activity in the Ala-triad point mutants of AtABC1K6.

## DISCUSSION

The study of the ABC1 family has been hampered by the previous inability to conclusively demonstrate catalytic function. Here, we report that an *A. thaliana* ortholog, AtABC1K6, possesses *in vitro* protein kinase activity requiring two modifications from standard *in vitro* assay buffers: i) provision of Mn^2+^ rather than Mg^2+^, and ii) low salt concentration. The requirement for Mn^2+^ suggests it, rather than Mg^2+^, is used as divalent cation co-factor. There is precedent for Mn^2+^-dependency among protein kinases, including a 2-component sensor kinase of the chloroplast stroma (33). The elimination of NaCl, on the other hand, is more surprising. *In vitro* kinase data from the literature corroborate an apparent salt sensitivity since the report of weak kinase activity from AtABC1K1 and AtABC1K3 used 50 mM NaCl (14, 15). This intermediate level of activity is consistent with the gradient of activity we report with our isolated plastoglobules (**Fig. 4A**).

Complete removal of buffer salt cannot be physiologically relevant to ABC1s, since the chloroplast stroma is thought to contain >100 mM KCl (34). However, the stroma can modulate ionic strength, primarily through transport of K^+^, which may impact ABC1 activity (35, 36). Further study will be necessary to determine whether physiological changes in stromal ionic strength can impact ABC1 activity *in vivo*. Regardless, we suggest that in manipulating ionic strength of the buffer we have identified a way to artificially induce regulatory rearrangements of the kinase structure that are a part of the natural regulatory cycle of AtABC1K6, leading to activation of kinase activity. All protein kinases are characterized by a spatially conserved arrangement of hydrophobic residues that create a pair of “spines” providing a strong but flexible scaffold (37–39). Rearrangements of these hydrophobic interactions are central to the structural dynamics that regulate activity, but are complex and specific to each kinase family. Reduced ionic strength in our buffer may influence the hydrophobic interactions within the AtABC1K6 spines, leading to activation of protein kinase activity.

An insert motif within the N-lobe called the KxGQ domain, unique to the ABC1 family, tightly packs over the active site cleft via hydrophobic interactions, occluding access of substrate (11). The placement of the KxGQ domain is comparable to the inactive conformations of ePKs, in which an Activation Loop is also folded over the active site cleft, preventing access of substrate (40, 41). In fact, two of the key distinguishing features of ABC1 proteins is their absence of an Activation Loop and the presence of the KxGQ domain. We thus propose that the KxGQ domain of ABC1 proteins is functionally analogous to the mobile Activation Loop of canonical ePKs, serving as an auto-regulatory feature that can expose or obscure the active site cleft to activate/inactivate the kinase. Moreover, the presence of the conserved Ala-triad adjacent to the ATP binding site, and the impact of mutating to Gly, highlights the importance of this adjoining region. In this work we show that such mutations greatly diminish kinase activity, which we suggest is due to a destabilization and unfolding of the β-sheet forming the ATP-binding pocket. However, further experimental and computational support is required to more fully explain this effect on kinase activity of AtABC1K6.

Our results with HsADCK3^NΔ254^ present a puzzle: why are the requirements for enzymatic activity that we demonstrate with AtABC1K6 – the salt sensitivity, the absolute dependency for Mn^2+^, or the significance of the intact Ala triad – not seen with HsADCK3^NΔ254^? The answer may lie in the extension sequence(s) flanking the conserved ABC1 kinase domain, which is removed on the N-terminal end from the assays with HsADCK3^NΔ254^, but left mostly intact with that of AtABC1K6^NΔ71^. In contrast to the highly conserved ABC1 domain, the extension sequences are highly variable in length and sequence, and often encode predicted lipid or protein binding motifs. Thus, the extensions may define the distinct functional identity of each ABC1 and regulate auto-inhibition that can be relieved only by a proper trigger. In this scenario, the absence of the N-terminal extension of HsADCK3^NΔ254^ may have weakened its auto-inhibitory capacity. Indeed, the crystal structure of the protein (PDB: 5I35) reveals multiple hallmarks of an active conformation, including DFG_in_, forming an Asp-His salt bridge, and αC-helix_in_, forming an Lys-Glu salt bridge (40, 41), indicating that the HsADCK3^NΔ254^ structure is primed for activation. Alternatively, unique catalytic or regulatory mechanisms may have emerged in each of the two proteins, reflecting their distinct functional roles and sub-cellular localizations.

Plastoglobule morphology is very dynamic during developmental transitions, which has suggested that they may play a role(s) in mediating development (42–46). Our demonstration that the plastoglobule-localized AtABC1K6 is necessary for the timely transition from vegetative to reproductive growth supports this notion and identifies a putative regulatory player. Although our assay with *abc1k6-1* plastoglobules and chloroplasts did not detect appreciable activity, AtABC1K6 may not be active at the developmental stage or condition from which our plastoglobules were isolated, or the AtABC1K6 activity is dwarfed by that of other plastoglobule ABC1 kinases. It remains unclear what substrates AtABC1K6 may target in affecting initiation of reproductive growth, but the biosynthetic enzymes of Jasmonic Acid (JA) biosynthesis that partially localize to the plastoglobule (4, 5, 47, 48), particularly AtLOX2 which our results suggest is remobilized to the plastoglobule in *abc1k6-1*, present an intriguing possibility. The phytohormone JA is a positive regulator of developmental senescence and tightly intertwined with the early steps in the switch to reproductive growth (23, 24, 49–55).

In sum, by modifying buffer conditions we have been able to demonstrate *in vitro* protein kinase activity with the plastoglobule-localized AtABC1K6. Our discovery of proper buffer conditions to facilitate protein kinase activity among AtABC1K6 and HsADCK3 presents the experimental means to explore the substrates, activity and regulation of this protein family, opening up a long-awaited opportunity to understand the functions of this enigmatic protein family.

## MATERIALS AND METHODS

### Plant material and growth conditions

The T-DNA insertion line SALK_129956C (*abc1k6-1*, At3g24190) was obtained from the Salk collection and homozygous mutants were screened using the following oligonucleotides: K6F: 5’-ATCGCGTATTGCAGGGGAT-3’, K6R: 5’-CGAGTTTAACAAAATCTTTTACTATG-3’ and LBb1 5’-GCGTGGACCGCTTGCTGCAACT-3’. Additionally, oligonucleotides for OE16 (At2g28900) were used as positive control for PCR reactions: OE16 F 5’-TGTTAGCACGCCGAAG-3’ and OE16 R 5’-CTTACCAACCGCTGAG-3’. Homozygous mutant plants were isolated using antibiotic selection on Murashige and Skoog agar media supplemented with kanamycin (50 μg/mL). For complementation, the coding sequence of AtABC1K6 was cloned into the vector pB7FWG2 under the control of the 35S promoter (56). The construct was introduced into *Agrobacterium tumefaciens* strain GV3101 and *abc1k6-1* mutants were transformed by floral dip (57). The T1 progeny were selected on soil by spraying BASTA herbicide one and two weeks after sowing. Homozygous plants for the transgene were selected based on their resistance to BASTA of the T3 progeny, which came from a T2 progeny with 75 % of resistance against the herbicide to ensure a single insertion event. If not indicated otherwise, *A. thaliana* plants were grown on soil under normal light conditions (LD, 16/8 h light/dark, 21ºC and 120 μE m^2^ s^−1^). Continuous light was applied using the same conditions but with a 24-hour light period.

### Measurement of photosynthetic parameters

*A. thaliana* wild-type and mutant plants grown under long day conditions for three weeks were dark adapted for 15 minutes and analyzed according to their chlorophyll a fluorescence as described in Espinoza-Corral et al. (2) using a pulse-modulated fluorometer (Imaging PAM and DUAL-PAM100; Walz).

### SDS-PAGE and immunoblot analyses

Total leaf or chloroplast sub-compartment fractions were separated on 12% SDS-polyacrylamide gels (58) and transferred to a nitrocellulose membrane (Amersham^™^, Protran^®^). The membrane was then incubated with monospecific polyclonal antisera (anti-FBN1a; AS06 116, anti-PsbA; AS05 084, anti-PsaF; AS06 104, anti-SBPase; AS15 2973, anti-LHC1a; AS01 005, anti-PsaD; AS09 461, anti-Cyt f; AS08 306, anti-PaO; AS11 1783 [Agrisera], anti-GFP; PA1 980A [Invitrogen], anti-His; TA150087 [OriGene]), followed by incubation with secondary polyclonal antisera HRP (Fisher scientific, 656120) and visualized by the enhanced chemiluminescence technique (GE Healthcare).

### A. thaliana protoplast isolation for confocal microscopy

Leaves from *A. thaliana* plants of 3 to 4 weeks old were excised and used for protoplast isolation following the method described by (59). Isolated protoplasts were then visualized under confocal microscopy.

### Chloroplast sub-compartment isolation

*A. thaliana* plants of 4 weeks old were used to isolated chloroplasts with subsequent separation of their sub-compartments as previously described in (5).

### Total protein leaf extraction

*A. thaliana* leaves were excised and ground in tubes containing glass beads. The ground tissue was resuspended in extraction buffer (50 mM Tris pH 7.5, 2% SDS and a protease inhibitor cocktail [P9599, Sigma]) and incubated for 30 minutes on ice. The samples were centrifuged for 30 minutes at 20.000 g and 4 ºC and the supernatant recovered containing the total protein extract. Protein content was measured using BCA method (Pierce™ BCA Protein Assay Kit, 23227, Thermo Scientific).

### *Heterologous expression of recombinant proteins in* E. coli

The coding sequence for the mature form of AtABC1K6 was cloned into the vector pMAL-c5x (New England BioLabs) as well as the point mutant versions of the protein. At the C-terminus of the protein an additional His-tag was added by cloning. These constructs were used to transform BL21 pLys cells which were induced for over-expression with 1 mM IPTG when they reached an OD_600_ of 0.6 - 0.8. The over-expression was performed over-night at 12ºC. After cell lysis by sonication in buffer containing 20 mM Tris-HCl pH 7.5 and 15 mM imidazole, the solution was centrifuged at 20,000g and 4 ºC for 30 min. The supernatant was transferred to equilibrated Ni^+2^–NTA beads (Thermo scientific) in 20 mM Tris-HCl pH 7.5 and 15 mM imidazole and incubated under rotation for 2 hours at 4ºC. Beads were pelleted by centrifugation at 100g and 4ºC for 2 minutes and then washed with buffer 20 mM Tris-HCl pH 7.5 and 15 mM imidazole for four times. The recombinant proteins were eluted by washing the beads with 20 mM Tris-HCl pH 7.5 and 500 mM imidazole. Eluted proteins were then dialyzed over night against kinase buffer (20 mM Tris-HCl pH 7.5, 5 mM MgCl_2_ and 0.5 mM MnCl_2_) and quantified by using Pierce™ BCA Protein Assay Kit (23227, Thermo scientific). In the case of protease digestions, Factor Xa (P8010S, New England BioLabs) or TEV protease (P8112S, New England BioLabs) were used for the cleavage of constructs in the backbone of pMAL-c5x and for His-MBP-[TEV]-ADCK3. The digestions were performed for one hour under room temperature for Factor Xa with the addition of 2 mM CaCl_2_ or one hour under 37°C for TEV protease with the addition of 0.1 mM DTT.

### Generation of point mutants

The constructs containing the AtABC1K6 gene for over-expression in *E. coli* using the vector pMAL5-x, as well as in *A. thaliana* using the vector pB7FWG2, were used as templates for the generation of point mutants. Point mutant variants were generated by inverse-PCR using the primers indicated in **SI Appendix, Dataset S4**. The inverse-PCR reaction was cleaned up and the template plasmid digested with DpnI for one hour at 37°C. After inactivating DpnI at 80°C for 10 min, the inverse-PCR product was ligated by adding T4 DNA ligase and T4 polynucleotide kinase (Thermo scientific). After bacterial transformation, the point mutants were confirmed by sequencing.

### In-vitro kinase assays

Heterologously expressed and purified recombinant protein or isolated plastoglobules were incubated with 3 μCi of [γ-^32^P] ATP in 40 μL kinase reaction buffer (20 mM Tris-HCl pH 7.5, 5 mM MgCl_2_ and 0.5 mM MnCl_2_). The reaction was incubated for 30 minutes to one hour (as described in the figure legend) at room temperature and stopped by adding 5 μL of SDS loading buffer without boiling the samples. The proteins were separated on 12% SDS-polyacrylamide gels followed by exposure to autoradiography films from overnight to up to one week depending on the signal intensity, as indicated in the figure legend.

### LC-MS/MS Proteomics and data analysis of heterologous MBP-AtABC1K6-His Protein

To delineate the boundaries of the MBP-AtABC1K6-His cleavage products, expressed and purified protein was re-suspended in SDS-PAGE sample buffer and heated at 60°C for 10 minutes. Samples were cooled and loaded onto a 12.5% pre-cast BioRad Criterion 1D gel and electrophoresed until the dye front began to run off the bottom of the gel. Electrophoresis was stopped and the gel fixed in 40% Methanol/20% Acetic Acid for 4 hours and stained with colloidal Coomassie Blue overnight. The gel was then de-stained using 10% Acetic Acid until the background was clear. Gel bands were digested in-gel according to Shevchenko, et. al. (60) with modifications. Briefly, gel bands were dehydrated using 100% acetonitrile and incubated with 10 mM dithiothreitol in 100 mM ammonium bicarbonate, pH ∼8, at 56°C for 45 min, dehydrated again and incubated in the dark with 50 mM iodoacetamide in 100 mM ammonium bicarbonate for 20min. Gel bands were then washed with ammonium bicarbonate and dehydrated again. Sequencing grade modified trypsin was prepared to 0.01 μg/μL in 50 mM ammonium bicarbonate and ∼100uL was added to each gel band so that the gel was completely submerged. Bands were then incubated at 37°C overnight. Peptides were extracted from the gel by water bath sonication in a solution of 60% Acetonitrile/1% Trifluoroacetic acid and vacuum dried to ∼2 μL. An injection of 5 μL was automatically made using a Thermo (www.thermo.com) EASYnLC 1000 onto a Thermo Acclaim PepMap RSLC 0.1mm x 20mm C18 trapping column and washed for ∼5 min with Buffer A. Bound peptides were then eluted over 35 min onto a Thermo Acclaim PepMap RSLC 0.075 mm x 250 mm resolving column with a gradient of 5% B to 40% B in 24 min, ramping to 90% B at 25 min and held at 90% B for the duration of the run (Buffer A = 99.9% Water/0.1% Formic Acid, Buffer B = 80% Acetonitrile/0.1% Formic Acid/19.9% Water) at a constant flow rate of 300 nL/min. Column temperature was maintained at a constant temperature of 50°C using an integrated column oven (PRSO-V1, Sonation GmbH, Biberach, Germany). Eluted peptides were sprayed into a ThermoScientific Q-Exactive mass spectrometer using a FlexSpray spray ion source. Survey scans were taken in the Orbitrap (70000 resolution, determined at m/z 200) and the top 15 ions in each survey scan were then subjected to automatic higher energy collision induced dissociation (HCD) with fragment spectra acquired at 17,500 resolution. The resulting MS/MS spectra were processed with the MaxQuant software program, version 1.6.17.0 (61). Peak lists were searched with the embedded Andromeda search engine (62) against the *Escherichia coli* UNIPROT database concatenated with the MBP-AtABC1K6-His heterologous protein sequence plus the common contaminant list appended by MaxQuant. Oxidation of methionine, deamidation of asparagine and glutamine, and N-terminal acetylation were set as variable modifications, carbamidomethylation was set as a fixed modification. Digestion mode was Trypsin/P with a maximum of 2 missed cleavages. MS/MS tolerance of the first search was 20 ppm, and main search was 4.5 ppm, with individualized peptide mass tolerance selected. False discovery rate (FDR) at peptide spectrum match and protein levels was set as 0.01. Filtering of resulting protein groups was performed manually at a fixed FDR of 0% by accepting protein IDs with the highest MaxQuant Scores until the first decoy protein ID was reached. The mass spectrometry proteomics data have been deposited to the ProteomeXchange Consortium via the PRIDE partner repository (https://www.ebi.ac.uk/pride/) in MIAPE-compliant format with the dataset identifier PXD027672.

### LC-MS/MS Proteomics and data analysis of plastoglobule samples

Lyophylized samples were re-suspended in 50 μL of 125 mM Tris, pH 6.8, 4% SDS, 20% glycerol, heated to 60°C for 5 min and electrophoresed into a 12% Tris-HCl Criterion gel from BioRad at 50V for ∼20 min or until the dye front moved just below the bottom of the sample wells. Electrophoresis was stopped and the gel stained with Coomassie Blue protein stain until bands appeared. Protein bands were excised from the gel and digested according to Shevchenko, et. al. (63) with modifications. Briefly, gel bands were dehydrated using 100% acetonitrile and incubated with 10mM dithiothreitol in 100 mM ammonium bicarbonate, pH∼8, at 56°C for 45 min, dehydrated again and incubated in the dark with 50 mM iodoacetamide in 100 mM ammonium bicarbonate for 20 min. Gel bands were then washed with ammonium bicarbonate and dehydrated again. Sequencing grade modified trypsin was prepared to 0.01 μg/μL in 50 mM ammonium bicarbonate and ∼100 uL of this was added to each gel band so that the gel was completely submerged. Bands were then incubated at 37°C overnight. Peptides were extracted from the gel by water bath sonication in a solution of 60% Acetonitrile (ACN) /1% Trifluoroacetic acid (TFA) and vacuum dried to ∼2 μL.

Injections of 5 μL were automatically made using a Thermo (www.thermo.com) EASYnLC 1200 onto a Thermo Acclaim PepMap RSLC 0.1mm x 20mm C18 trapping column and washed for ∼5 min with Buffer A. Bound peptides were then eluted over 65 min onto a Thermo Acclaim PepMap RSLC 0.075 mm x 500 mm resolving column with a gradient of 8% B to 40% B in 54 min, ramping to 90% B at 55 min and held at 90% B for the duration of the run (Buffer A = 99.9% Water/0.1% Formic Acid, Buffer B = 80% Acetonitrile/0.1% Formic Acid/19.9% Water) at a constant flow rate of 300 nL/min. Column temperature was maintained at a constant temperature of 50°C using and integrated column oven (PRSO-V2, Sonation GmbH, Biberach, Germany).

Eluted peptides were sprayed into a ThermoScientific Q-Exactive HF-X mass spectrometer (www.thermo.com) using a FlexSpray spray ion source. Survey scans were taken in the Orbitrap (60000 resolution, determined at m/z 200) and the top 15 ions in each survey scan are then subjected to automatic higher energy collision induced dissociation (HCD) with fragment spectra acquired at 7,500 resolution. The resulting MS/MS spectra were converted to peak lists using MaxQuant, v1.6.17.0, and searched against a database containing all *A. thaliana* protein sequences available from TAIR10 (downloaded from www.arabidopsis.org), with common laboratory contaminants using the Andromeda search algorithm, a part of the MaxQuant environment. The mass spectrometry proteomics data have been deposited to the ProteomeXchange Consortium via the PRIDE partner repository (https://www.ebi.ac.uk/pride/) in MIAPE-compliant format with the dataset identifier PXD028493.

### LC-MS/MS Proteomics and data analysis of leaf tissue samples

The samples (∼100 μg each) were precipitated using chloroform:methanol (1:4) to remove buffer salts and SDS. Protein pellets were then re-suspended in 100mM Tris-HCl, pH 8.0, supplemented with 4% (w/v) sodium deocycholate (SDC) and digested according to Humphrey, et al. (64). Following digestion, SDC was removed by phase extraction, peptides desalted using C18 StageTips and eluates dried by vacuum centrifugation. Each sample was then re-suspended in 20 μL of 2% acetonitrile/0.1% trifluoroacetic acid.

Injections of ∼2 μg were automatically made using a Thermo (www.thermo.com) EASYnLC 1200 onto a Thermo Acclaim PepMap RSLC 0.1 mm x 20 mm C18 trapping column and washed for ∼5min with Buffer A. Bound peptides were then eluted over 125 min onto a Thermo Acclaim PepMap RSLC 0.075 mm x 500 mm resolving column with a gradient of 8% B to 40% B in 114 min, ramping to 90% B at 115 min and held at 90% B for the duration of the run (Buffer A = 99.9% Water/0.1% Formic Acid, Buffer B = 80% Acetonitrile/0.1% Formic Acid/19.9% Water) at a constant flow rate of 300 nL/min. Column temperature was maintained at a constant temperature of 50°C using and integrated column oven (PRSO-V2, Sonation GmbH, Biberach, Germany).

Eluted peptides were sprayed into a ThermoScientific Q-Exactive HF-X mass spectrometer (www.thermo.com) using a FlexSpray spray ion source. Survey scans were taken in the Orbitrap (35000 resolution, determined at m/z 200) and the top 15 ions in each survey scan are then subjected to automatic higher energy collision induced dissociation (HCD) with fragment spectra acquired at 7,500 resolution. The resulting MS/MS spectra were converted to peak lists using MaxQuant, v1.6.17.0, and searched against a database containing all *A. thaliana* protein sequences available from TAIR10 (downloaded from www.arabidopsis.org), with common laboratory contaminants using the Andromeda search algorithm, a part of the MaxQuant environment. The mass spectrometry proteomics data have been deposited to the ProteomeXchange Consortium via the PRIDE partner repository (https://www.ebi.ac.uk/pride/) in MIAPE-compliant format with the dataset identifier PXD028277.

## ACKNOWLEDGEMENTS

This work was supported by the USDA Umbrella Program (MICL02633), AgBioResearch startup funds, and Michigan State University startup funds to P.K.L. We thank Dr. David Pagliarini (Washington University of St. Louis) for donation of HsADCK3 constructs, Dr. Alsonso I. Carvajal (Ludwig Maximilians University Munich) for help on bioinformatics analyses, Dr. Serena Schwenkert (Ludwig Maximilians University Munich) for providing the DUAL-PAM equipment for chlorophyll fluorescence measurements and the anti-LHC antibodies, Dr. John Wang (Michigan State University) for providing the anti-MBP antibody, Dr. John Froehlich (Michigan State University) for providing the anti-ATP synthase antibody, and Drs. Dean Della Penna and Christoph Benning (Michigan State University) for valuable feedback on the manuscript.

