## Supplemental Figures for "The plastoglobule-localized AtABC1K6 is a Mn^2+^-dependent protein kinase necessary for timely transition to reproductive growth"

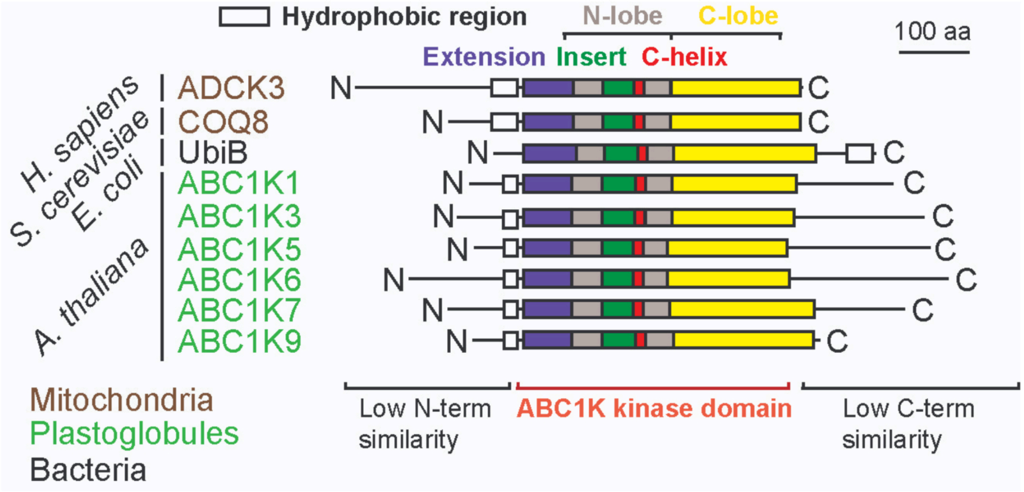


**Supplemental Figure 1.** Sequence structure of the ABC1 kinase family. Selected ABC1 orthologs from human, yeast, and bacteria, along with the six plastoglobule-localized ABC1 orthologs of *A. thaliana*, are aligned to highlight the conservation of the ABC1 kinase domain. Note the N- and C-terminal extensions outside the ABC1 domain which are highly divergent in length and sequence. The classical bi-lobed structure of eukaryotic protein kinases (ePKs) is seen in the ABC1 domain (grey and yellow). However, a unique insert (green) is found within the N-lobe that includes the KxGQ motif and is found lying over the substrate binding pocket in the HsADCK3$^{N\Delta254}$ crystal structure. It is this insert that we propose to be the functional analog of the Regulatory Domain found within ePKs.

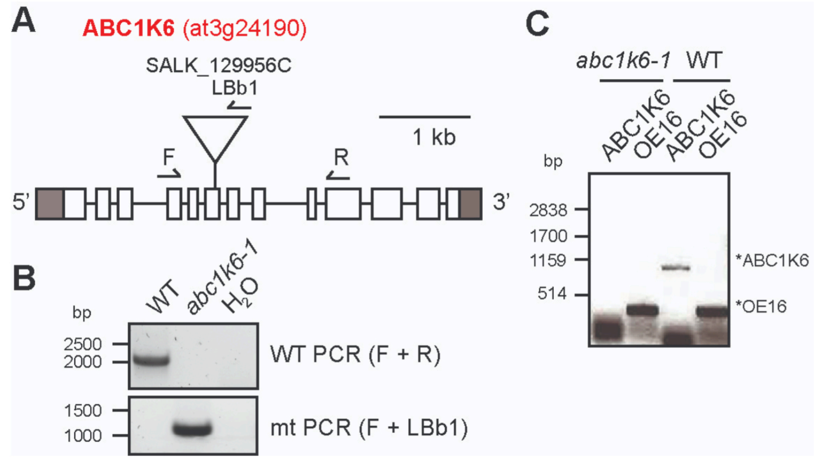


**Supplemental Figure 2. Genotyping of *abc1k6-1*. A,** gene structure of abc1k6 (at3g24190), the position of the T-DNA insertion of *abc1k6-1* and primers for genotyping. **B,** PCR amplification of genomic DNA confirming the homozygosity of the isolated *abc1k6-1* line. **C,** RT-PCR of leaf tissue cDNA indicating no accumulation of full length transcript.

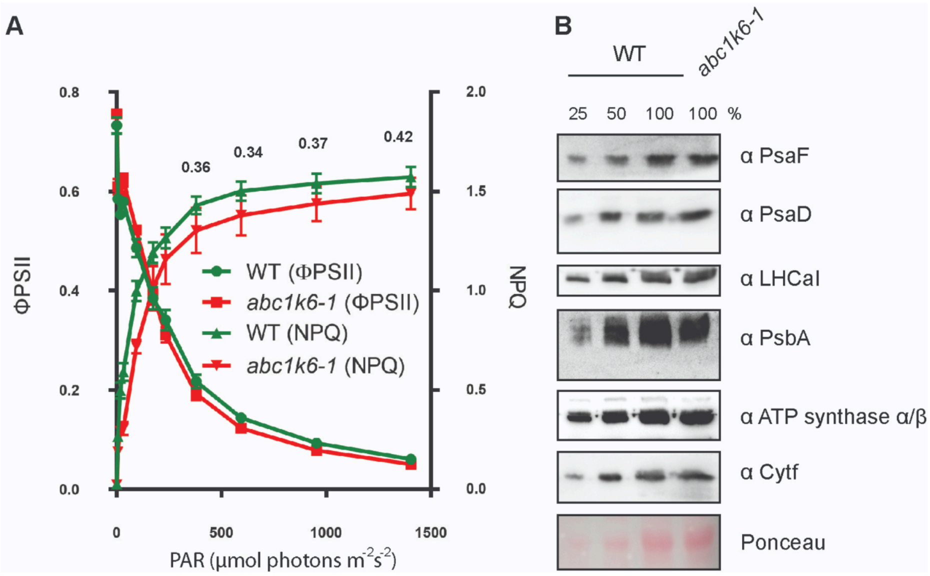


**Supplemental Figure 3. Photosynthetic phenotype of *abc1k6-1*.** **A**, Quantum yield of Photosystem II (ΦPSII) and non-photochemical quenching (NPQ) plotted as a light-response curve. No statistically significant differences were observed between genotypes. Student's t-test p-values are displayed above the NPQ values of the four highest light intensities. **B,** Immunoblot of total leaf samples from Col-0 (WT) and abc1k6-1 illustrating the equal abundance of subunits of PSII, PSI, LHCI, cyt b6f, and ATP Synthase complexes.

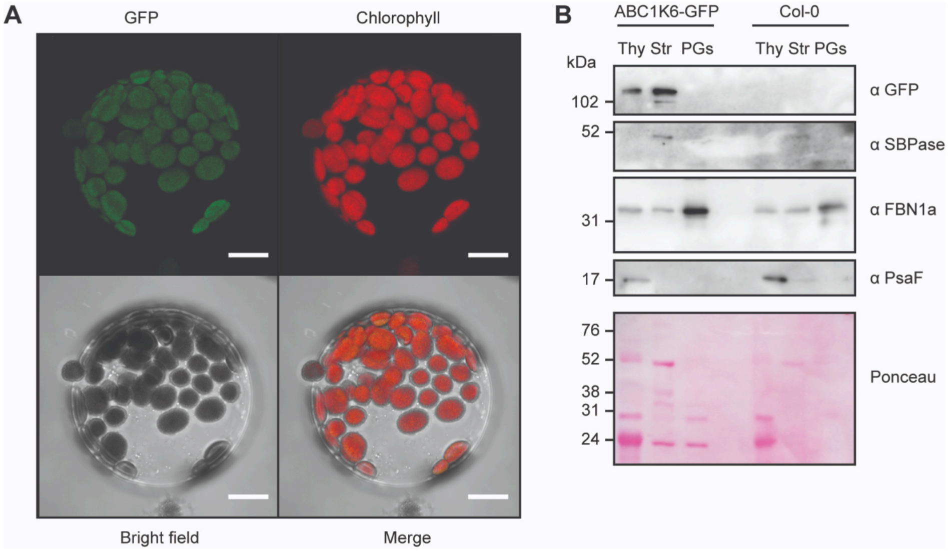


**Supplemental Figure 4.** Biochemical and Microscopic Investigation of the Sub-ecllular Loaclaization of ABC1K6-GFP in Comp3. **A,** Isolated protoplasts from the Comp3 line were imaged by confocal microscopy. Scale bar is 10 µm. **B,** Immunoblots of chloroplast sub-compartments from Comp3 (ABC1K6-GFP) and wild-type (Col-0). Anti-SBPase is a marker for stroma (str), anti-FBN1a for plastoglobules (PGs), and anti-PsaF for thylakoid (thy).

A

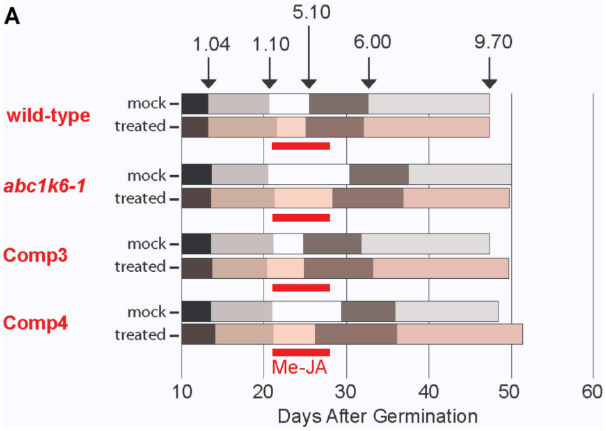


B

wild-type *abc1k6-1* Comp3 Comp4

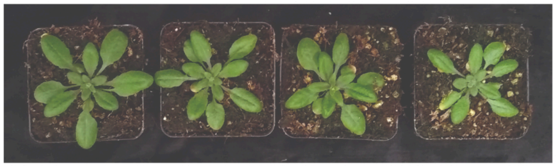

**Supplemental Figure 5. Me-JA treatment Fails to Complement the developmental Phenotype of *abc1k6-1*. A,** schematic of developmental rate of each genotype under mock and methyl-jasmonic acid (Me-JA) treatments. The time period within which plants were sprayed daily with water (mock) or Me-JA is indicated with the red underline, from 21 to 28 days after germination. No statistically significant differences ($p$-value $< 0.05$) were observed when comparing mock to Me-JA treatment within each genotype. n = 10 individual plants. **B,** Images of representative plants at 4 weeks of age.

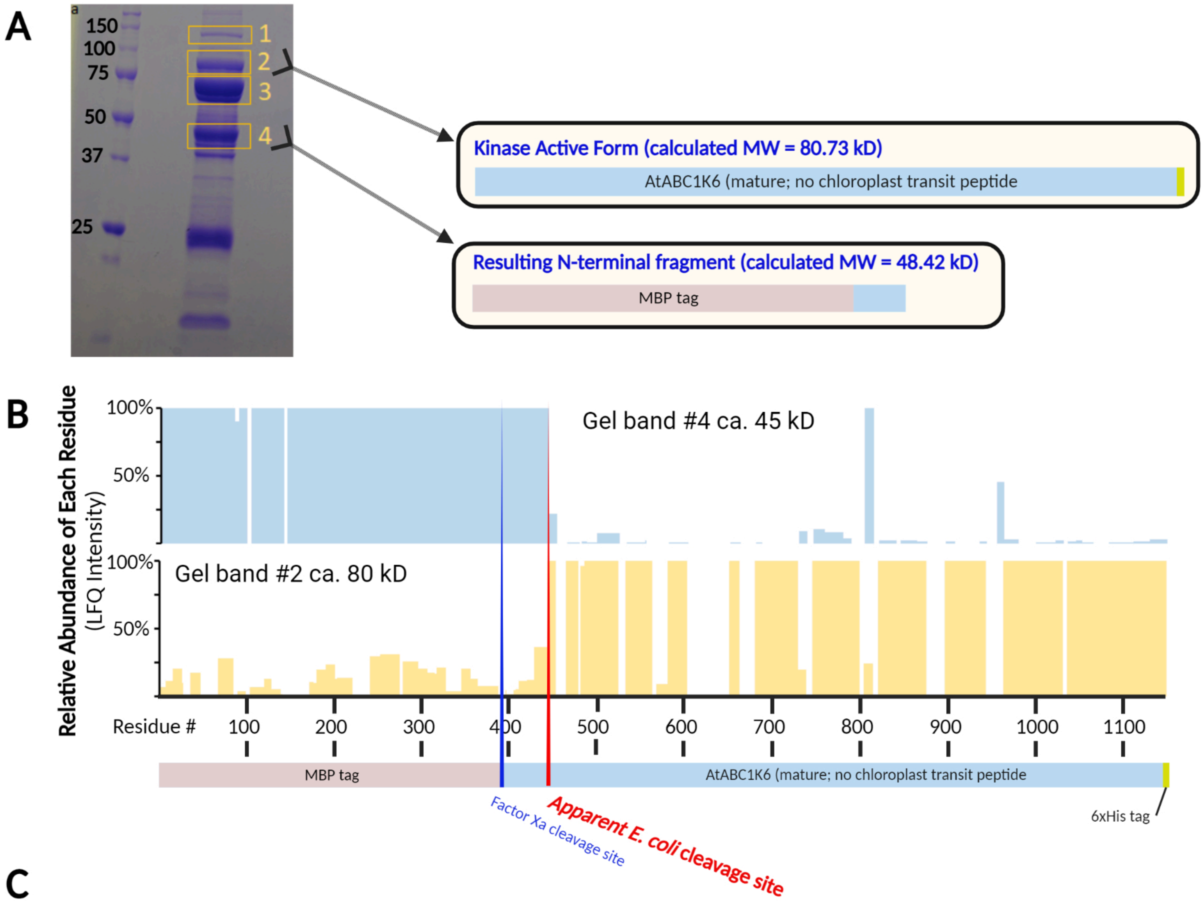


C

MKIEEGKLVIWINGDKGYNGLAEVGKKFEKDTGIKVTVEHPDKLEEKFPQVAATGDGPDIIFWAHDRFGGYAQSGLLAEITPDKAFQDKLYPFTWDAVRY
NGKLIAYPIAVEALSLIYNKDLLPNPPKTWEEIPALDKELKAKGKSALMFNLQEPYFTWPLIAADGGYAFKYENGKYDIKDVGVDNAGAKAGLTFLVDLI
KNKHMNADTDYSIAEAAFNKGETAMTINGPWAWSNIDTSKVNYGVTVLPTFKGQPSKPFVGVLSAGINAASPNKELAKEFLENYLLTDEGLEAVNKDKPL
GAVALKSYEEELVKDPRIAATMENAQKGEIMPNIPQMSAFWYAVRTAVINAASGRQTVDEALKDAQTNSSSNNNNNNNNNNLGIEGRISHMVSSRRSFVS
GIPHRNKRSRQILAVATDPKPTQTSPPKSTTVNGSSSPSSASKVVNNNVSTRINDVSKEIKRVRAQMEEDEQLSVLMRGLRGQNLKDSVFADDNIQLRLV
ETGESSEFLPLVYDPETISAYWGKRPRAVASRVIQLLSVAGGFLSRIAGDVINKKVKENEVARAIELREIVTSLGPAYIKLGQALSIRPDILSPAAMTEL
QKLCDKVPSYPDDVAMALIEEELGKPWYDVYSELSPSPIAAASLGQVYKGRLKENGDLVAVKVQRPFVLETVTVDLFVIRNLGLFLRKFPQVSVDVVGLV
DEWAARFFEELDYVNEGENGTYFAEMMKKDLPQVVVPKTYQKYTSRKVLTTSWIDGEKLSQSIESDVGELVNVGVICYLKQLLDTGFFHADPHPGNMIRT
PDGKLAILDFGLVTKLTDDQKYGMIEAIAHLIHRDYDAIVKDFVKLGFIPDGVNLAPILPVLAKVFDQALEGGGAKNINFQELAADLAQITFDYPFRIPP
YFALIIRAIGVLEGIALVGNPEFAIVDEAYPYIAQRLLTDESPRLREALRYTIYGKTGVFDAERFIDVMQAFETFITAAKSGGGEDMNGGMAEMALMQSK
TSSLVPMFPASASQPDQPVQTRVALSFLLSEKGNFFREFLLDEIVKGIDAITREQLVQAMAVFGFRNATPIFGMLPPTLGPFKPAALLPSVTEEDKVILN
NVQKVIEFLTARSSMSNNPDQVVDVSQVVRELLPVLPGISATVLPEILSRLGSRVMARIVRDAFLHHHHHH

**Supplemental Figure 6. Mass Spectrometry-based Proteomics of Excised Gel Bands Identifies Proteolytic Cleavage following Residue K443 of the MBP-AtABC1K6-His Protein. A,** The mature form of AtABC1K6 (i.e. lacking the putative chloroplast transit peptide which is removed and degraded following chloroplast import) was cloned in-frame with an N-terminal Maltose Binding Protein (MBP) and a C-terminal 6xHis tag. After heterologous expression in *Escherichia coli* and purification using Ni-NTA beads, the protein sample was separated by SDS-PAGE, gel bands excised, and subjected to quantitative LC-MS/MS analysis to determine the identitiy of each protein sequence. **B,** Each residue's relative abundance, normalized to its highest abundance across the four gel bands, is plotted. A value of 100% indicates the residue is most abundant in the respective gel band. The results indicate the a cleavage event between residue K443 and V444 that results in a protein product of ca. 80 kD comprising most of the mature AtABC1K6 protein sequence that coincides with the protein size capable of in vitro autophosphorylation. **C,** Sequence of the full-length recombinant protein. The apparent cleaveage site occuring during *E. coli* expression or purification is highlighted in red underline.

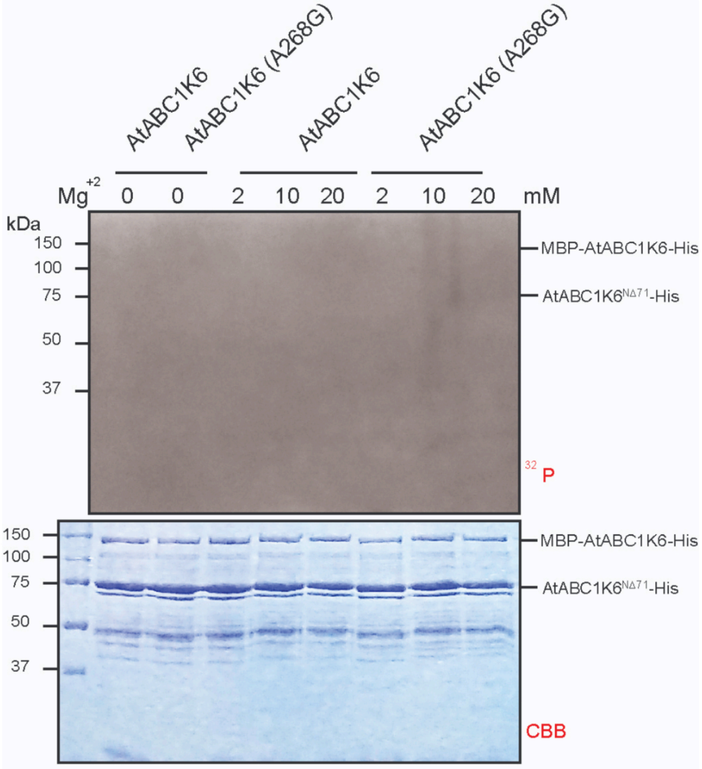


**Supplemental Figure 7.** Heterologously Expressed and Purified AtABC1K6 Lacks Kinase Activity in the Presence of $Mg^{2+}$ Alone. The recombinant proteins, MBP-ABC1K6-His wild type and its point mutant A268G, were heterologusly expressed in *E. coli* and purify by $Ni^{+2}$-NTA beads. In vitro kinase assays were performed by supplementing increasing concentrations of $Mg^{2+}$ into the reaction buffer containing the same amount of radiolabeled ATP. After separating two µg protein per lane on an SDS-PAGE gel, the gel was dried and exposed to an autoradiography film for one week.
